# Dual-stream cortical feedbacks mediate sensory prediction

**DOI:** 10.1101/2022.07.12.499695

**Authors:** Qian Chu, Ou Ma, Yuqi Hang, Xing Tian

## Abstract

Predictions are constantly generated from diverse sources to optimize cognitive functions in the ever-changing environment. However, the neural origin and generation process of top-down induced prediction remain elusive. We hypothesized that motor-based and memory-based predictions are mediated by distinct feedback networks from motor and memory systems to the sensory cortices. Using fMRI and a dual imagery paradigm, we showed that motor and memory upstream systems excited the auditory cortex in a content-specific manner. Moreover, the inferior and posterior parts of the parietal lobe differentially relayed predictive signals in motor-to-sensory and memory-to-sensory networks. Our results reveal the functionally distinct neural networks that mediate top-down sensory prediction and ground the neurocomputational basis of predictive processing.

## Introduction

Generating predictions is a trait of adaptive organisms to efficiently interact with the environment (Conant and Ashby, 1970; Friston, 2010; Schultz et al., 1997). For example, a seminal trend in cognitive neuroscience considers perception to depend on dynamic predictions based on the internal models of external world (Bar, 2007; de Lange et al., 2018; Rao and Ballard, 1999). In contrast to the *feedforward* information flow from sensory to non-sensory areas, coordinated *feedback* projections from non-sensory to sensory areas provide a neural substrate for conveying top-down sensory predictions (Keller and Mrsic-Flogel, 2018).

How feedback projections convey predictive signals in the human brain remains enigmatic. Theoretically, the action-perception loop that links an agent’s cognitive system and the environment necessitates multiple forms of predictions. One category of predictions is motor based. According to theories of motor control, the agent could use a copy of the endogenous motor command and a model of action-consequence coupling to predict the sensory consequences of actions (McNamee and Wolpert, 2019; Shadmehr et al., 2010; Wolpert and Ghahramani, 2000). Motor-based predictions could be used for world state estimation (Wolpert et al., 1995). The resulting prediction error could drive immediate motor correction as well as long-term motor learning (Jordan and Keller, 2020). Whereas, predictions that do not involve an agent’s actions are exemplified by the suppression of neural response to statistically organized stimuli (e.g., structured sequences (Garrido et al., 2009; Todorovic et al., 2011) or associated pairs (Garner and Keller, 2022; Kok et al., 2012)). Humans learn rich statistical regularities in the external world and utilize the exogenous information by transforming memory traces into sensory predictions. The combination of motor-based and memory-based predictive algorithms constructs a dual-stream prediction model (DSPM) (Tian and Poeppel, 2013; Tian et al., 2016) – motor and memory systems could reverse their traditionally assumed roles as receivers of sensory information to act as independent sources that provide endogenous and exogenous information for generating sensory prediction (Figure 1a).

**Figure 1:**
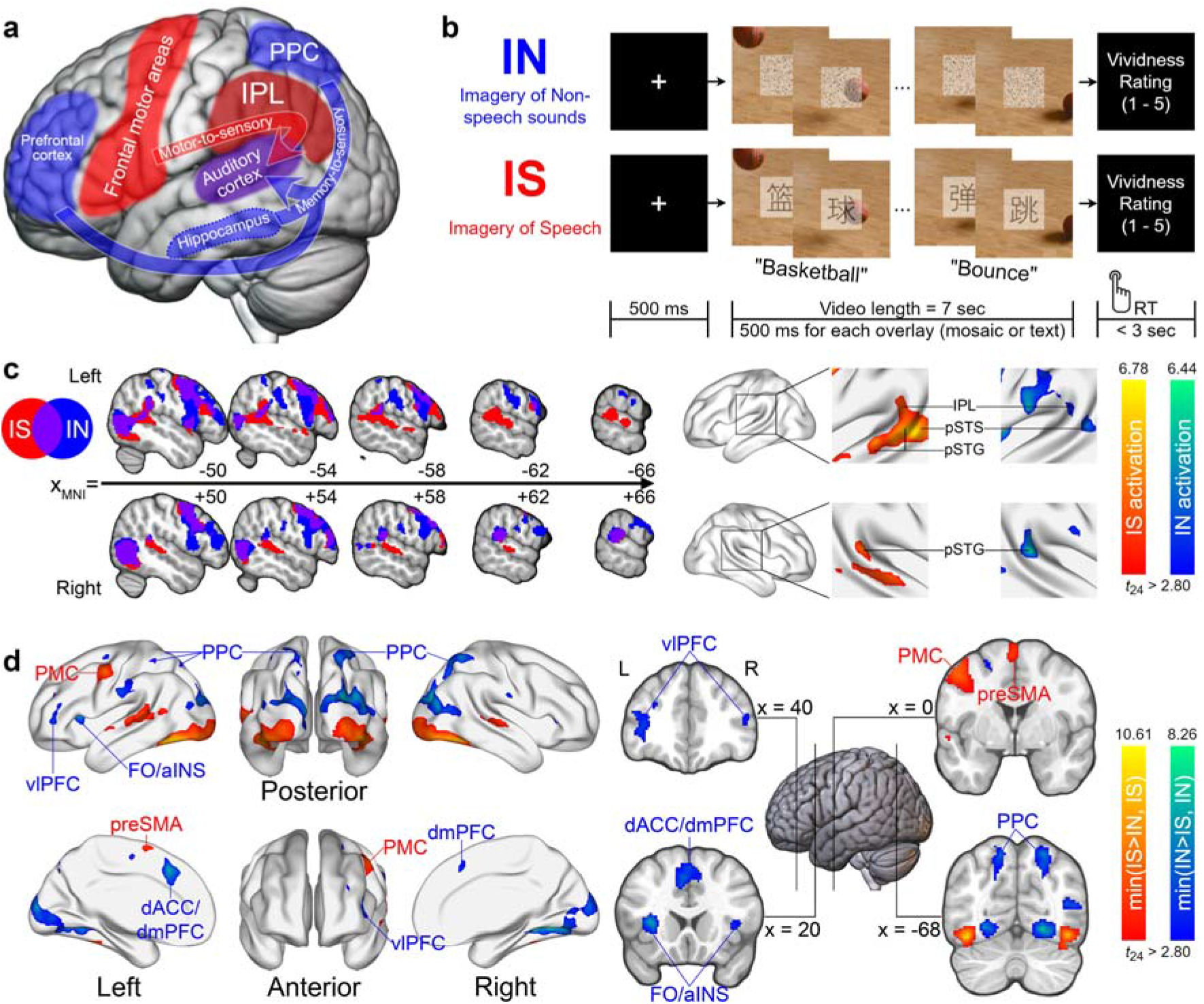
Model of distinct pathways for generating prediction, experimental paradigms, and fMRI results of univariate analyses. (**a**) The Dual-Stream Prediction Model (DSPM). The model posits that auditory representations in the temporal area can be established by two feedback streams. The motor-to-sensory stream originates from the frontal motor network where speech plan encoding is carried out. A copy of the motor plan (efference copy), relaying via the inferior parietal lobe, establishes auditory representations in the auditory cortex to predict the sensory consequence of speech action. The memory-to-sensory stream, originating in a distributed memory network including the prefrontal cortex, hippocampus, and superior parietal lobe reconstructs auditory representations in the auditory system via memory retrieval. (**b**) Experimental paradigm. Following a 500 ms fixation period, participants watched a muted video of objects in motion (frames from the bouncing basketball video are used for illustration). Participants were asked to imagine sounds ought to be in the video (e.g., the whomp of a basketball hitting the floor repeatedly) in the *IN* condition and imagine saying characters superimposed on the video in the *IS* condition. (**c**) Activations in the inferior parietal and superior temporal regions during *IS* and *IN*. Top: activations in the left hemisphere. Bottom: activations in the right hemisphere. Left: the mosaic view. Colored voxels were activated significantly in *IS* (red), *IN* (blue), or both (purple). Right: Thresholded surface rendering with *t*-value indicated by the color bar. See also Figure S1. (**d**)Thresholded surface rendering showing the conjunctions (minimum statistic) between 1) *IS* > *IN* and *IS*, and 2) *IN* > *IS* and *IN*. *IS* induced stronger activations in the left PMC and preSMA, whereas *IN* induced stronger activations in the bilateral fronto-parietal and cingulo-opercular networks.

Methodological challenges also obstruct the investigation of the neural basis of prediction. This is partly because of the spatial-temporal overlapping between feedback prediction and feedforward input during perception (Keller and Mrsic-Flogel, 2018). Moreover, most studies investigate predictive processing by probing how prediction modulates perception, granting them only indirect access to feedback predictive signals. This indirect modulation approach that focuses on the functions of prediction is hard, if not impossible to reveal the neural origin and generation processes that constrain the cognitive computations and the neural implementation of predictive processing from a system perspective.

Mental imagery serves as a promising paradigm for directly scrutinizing what and how feedback projections convey predictive signals. Imagery, a cognitive capacity to endogenously create episodic mental states (Langland-Hassan, 2020), has been widely reported to elicit perceptual-like neural representations (Bunzeck et al., 2005; Hubbard, 2010; Kosslyn et al., 1999; Kraemer et al., 2005; O’Craven and Kanwisher, 2000; Zatorre et al., 1996) resulted from top-down connectivity (Dentico et al., 2014; Dijkstra et al., 2017; Pearson, 2019). Because sensory prediction requires activating similar sensory representations of possible outcomes as imagery, imagery has been argued to be a mental realization of prediction (Moulton and Kosslyn, 2009) and exploit the same set of internal models as implemented in predictive processing (Langland-Hassan, 2016; Williams, 2021). Consistent with this proposal, mental imagery suppresses perceptual responses, similar as prediction does (Kilteni et al., 2018; Tian et al., 2018).

Therefore, we leveraged mental imagery to investigate feedback projections that establish auditory representations in the absence of confounding feedforward signals so as to trace the neural origin of predictions. Moreover, our novel dual-imagery paradigm maximized the differences between motor-based and memory-based prediction as participants were asked to imagining speech or natural sounds that human articulators cannot produce (Figure 1b). The DSPM model and preliminary empirical findings (Li et al., 2020; Ma and Tian, 2019; Tian and Poeppel, 2010; Tian *et al*., 2016) derive three major experimental predictions. First, both motor-based and memory-based predictions in different types of imagery would reactivate the auditory cortex without external acoustic stimulation. Second, the upstream networks for generating sensory predictions should be distinct. Motor-based imagery would activate the frontal motor network, whereas memory-based imagery would involve the frontal-parietal and hippocampal networks. Third and most importantly, information would flow directionally from motor or memory upstream systems to auditory areas in distinct functional feedback networks that mediate the generation of prediction. The parietal lobe in particular would relay feedback projections, with posterior parietal cortex (PPC) subserving memory-based prediction (Dijkstra *et al*., 2017; Sestieri et al., 2017) and inferior parietal lobe (IPL) as a sensorimotor interface in speech (Hickok, 2012; Hickok and Poeppel, 2007) subserving motor-based prediction (Li *et a*. 2020; Tian and Poeppel, 2010; Tian *et al*., 2016). By using multiple analyses in conjunction, we obtained evidence that supported our hypotheses and revealed the origin, structure, and endpoint of feedback connections in generating predictions.

## Results

Participants (*N* = 25) were familiarized with ten categories of 7-second videos featuring moving objects moving objects making sounds (e.g., a bouncing basketball, exploding firecrackers, a flowing stream, etc.). The videos were then muted in the later imagery sessions (Figure 1b). Participants were instructed to recall sounds that were present in the video (Imagery of Non-speech sounds,*IN*) or imagine speaking sentences (*Imagery of Speech, IS*) describing the scenes according to visual character cues superimposed on the center of the video (e.g.,“A basketball bounces on the wooden floor over and over”; see Table S1 for sentences describing all 10 videos). In *IN*, comparable square mosaics were overlaid, keeping the net visual intensity consistent across *IS* and *IN* (Methods). The *IN* sessions preceded *IS* sessions such that participants were unaware of linguistic descriptions of the video and thus minimizing the possibility of their engaging imagery of speech during *IN*. After each trial, participants rated the vividness of imagery (range = 1 - 5). The length of the video yielded a long stimulus duration that increases the signal-to-noise ratio of imagery-related neural activities, and the dual imagery paradigm optimally bifurcates the motor and memory sources of auditory prediction in the same visual context. Two hearing conditions that were comparable to *IS* and *IN* were also included to locate auditory representations (Methods).

Throughout the paper, we define significance in whole-brain analyses as voxel-wise *P* < 0.005 and cluster-wise *P*_FDR_ < 0.05. We are refrained from discussing effects in the occipital lobe since they were resulted from visual stimulation.

### Behavioral results

The completion and success of mental imagery are hard to assess behaviorally because imagery is an internal experience. We relied on the timeliness of the participants’ vividness report to infer whether they performed the imagery tasks instructed. Participants actively engaged in the imagery as the response rate of vividness rating after each trial was at ceiling (mean = 98.20%), with higher vividness ratings in *IS* than *IN* (*t*_24_ = 5.57,*P* = 10^-5^, two-sided paired *t*-test).

### Common activations in auditory cortices accompanied by differential motor and memory activations

To test the hypothesis that the auditory cortex is activated as the sensory endpoints of feedback signaling, we first carried out whole-brain univariate analyses. We found overlapping activations in both *IS* and *IN* in the bilateral posterior part of superior temporal gyri and sulci (pSTG and pSTS). The common activation in both *IS* and *IN* also extended to the left inferior parietal lobe (IPL) that anatomically covered parts of parietal operculum, posterior supramarginal gyrus, and planum temporale (Figure 1c, for whole-brain surface rendering see Figure S1). In *IS*, activations also extended to left anterior STG (aSTG), consistent with previous findings of aSTG harboring higher-level linguistic representations (e.g., phonemes and words (DeWitt and Rauschecker, 2012)). Activations at pSTG and IPL were observed in the hearing conditions (Figure S2), further supporting that these regions mediate auditory-like representations.

Next, we contrasted *IS* with *IN* to examine differential activations that would likely distinguish upstream networks underlying prediction generation (Figure 1d). We took a minimum statistics approach (Nichols et al., 2005) to select voxels that showed both significant activity during one type of imagery and significant difference over the other (e.g., *IS* > *IN* masked with *IS* activations). *IS* induced stronger effects than *IN* in the frontal motor network, including the left premotor cortex (PMC) and pre-supplementary motor area (preSMA). *IN* activated the frontoparietal network (FPN) comprising the left ventrolateral prefrontal cortex (vlPFC) and bilateral posterior parietal cortex (PPC), and the cingulo-opercular network (CON) comprising the dorsal anterior cingulate cortex (dACC) and bilateral frontal operculum/anterior insular (FO/aINS).

### Motor, memory, and auditory systems represent imagery contents

To further test whether the activated areas represent imagery contents, we trained support vector machine-based classifiers to decode the imagery associated with 10 video categories in *IS* and *IN* (multivoxel pattern analysis, MVPA). To efficiently evaluate decoding accuracy across the brain, we conducted leave-one-run-out searchlight analyses with varying spherical radii ranging from 1 to 8 voxels (Methods).

High decoding accuracy observed in the visual cortex demonstrated the validity of our decoding method since the videos differed in visual stimulation. Moreover, we found above-chance accuracy (chance level = 10%) in bilateral pSTG and left IPL in both *IS* (Figure 2a), *IN* (Figure 2b), and comparable hearing conditions (Figure S3). These results support our hypothesis that specific auditory representations were activated in a top-down manner as auditory endpoints in the feedback networks.

**Figure 2:**
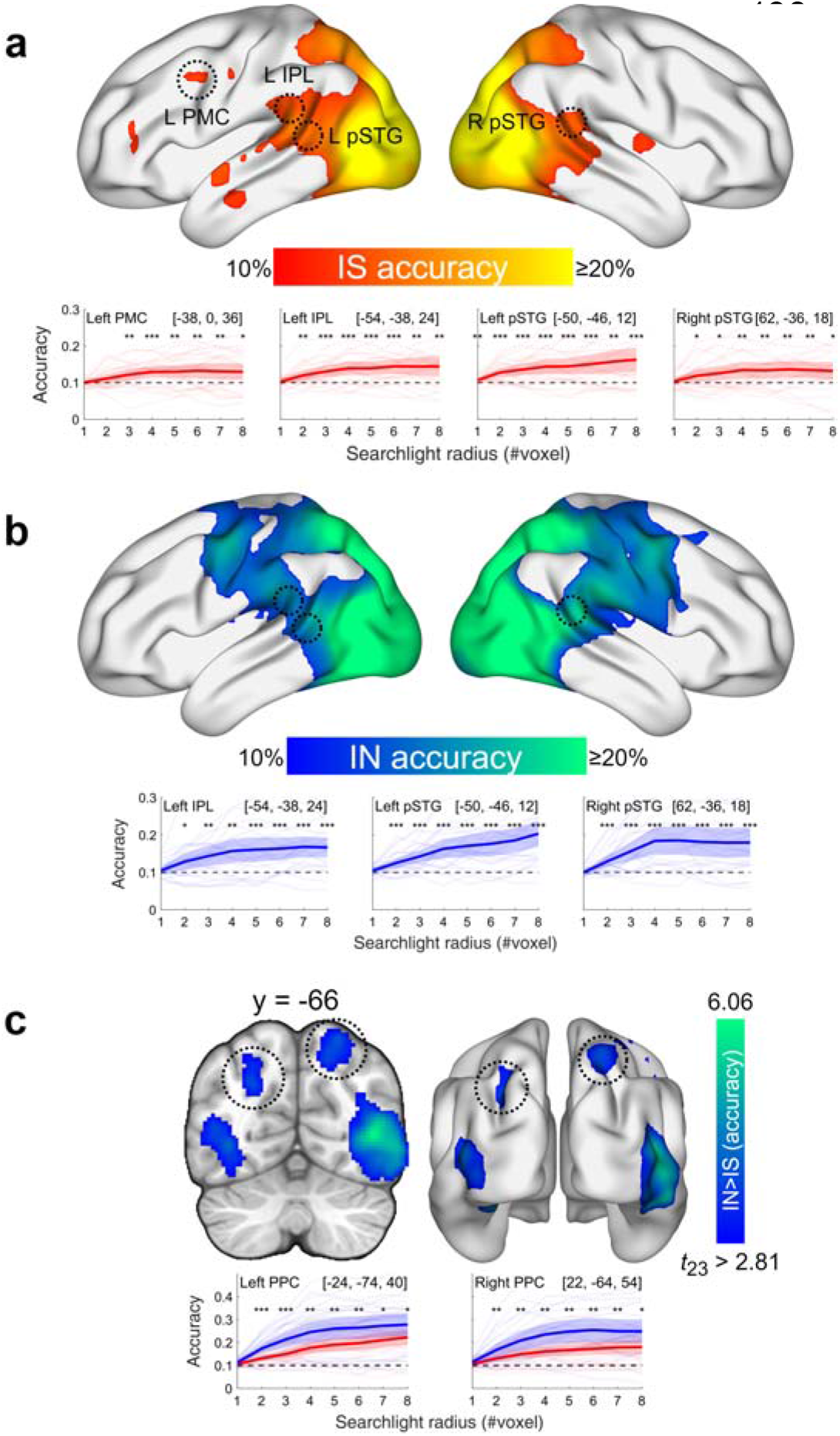
Results of multivoxel pattern analysis. (**a**)Decoding of video categories in *IS*. Top: thresholded surface rendering of decoding accuracy using a moving searchlight with a radius of 4 voxels. Bottom: decoding accuracy at regions of interest across different radii (1 – 8 voxels). The triplet numbers in brackets denote MNI coordinate of the searchlight center. Asterisks denote significance level of decoding accuracy above chance level (10%). (**b**) Similar to (a) but for classification in *IN*. (**c**) Top: a coronal view and a surface rendering of areas showing higher decoding accuracy in *IN* than *IS*. Bottom: classifier performance in bilateral PPC during *IS* and *IN* across searchlight radii. Asterisks denote significance level of decoding accuracy higher in *IN* than *IS*. For all panels, error bars indicate 95% confidence interval. \**P* < 0.05; \*\**P* < 0.01; \*\*\**P* < 0.001.

Consistent with univariate results, significant decoding of videos was found in the left PMC in *IS*. This decoding of imagery contents in the frontal motor region without participants’ overt movement suggests a motor representation space in the motor upstream network (Figure 2a).

For *IN*, decoding accuracy was significantly above chance in bilateral PPC but not in vlPFC nor in the cingulo-opercular network (Figure 2b). Despite significant decoding observed in bilateral PPC during *IS*, two-sided paired *t*-tests showed that the decoding accuracy in parts of PPC (left intraparietal sulcus and right superior parietal lobule) was significantly higher in *IN* than that in *IS* reliably across searchlight radii (Figure 2c), suggesting memory representations in PPC in addition to putatively visual representations commonly available in both conditions (confirmed by a cross-classification analysis, Figure S4).

Putting together the univariate and MVPA results, the selective activations and content specificity of PMC in *IS*, PPC in *IN*, and the auditory cortex in both conditions supported our first hypothesis of common sensory endpoint and our second hypothesis of differential upstream systems for motor-based and memory-based prediction. We next tested our last hypothesis about the feedback structures mediating the two types of predictions by examining the cortico-cortical connectivity with dynamic causal modeling (DCM).

### Motor-to-sensory and memory-to-sensory networks assessed by dynamic causal modeling

For connectivity analyses, we selected regions of interest (ROIs) based on univariate and MVPA results. The representative voxel coordinate of each ROI and their associated *t*-values for each contrast are reported in Table S2 and all selected voxels are visualized in Figure 3a. Our criteria are summarized here. For auditory ROIs, we selected areas that showed increased BOLD magnitude and representational patterns during both *IS* and *IN*, leading to our choice of left pSTG (sphere center *x* = −50, *y* = −46, *z* = 12) and its right homologue (sphere center *x* = 62, *y* = −36, *z* = 18). Given its consistent appearance revealed by multiple analyses, left IPL (sphere center *x* = −54, *y* = −38, *z* = 24) was also selected to test whether it serves as a mediating hub for motor-to-sensory and memory-to-sensory feedback networks. As for the motor ROI, we included left PMC (sphere center *x* = −38, *y* = 0, *z* = 36) based on its significantly higher activity during *IS* than IN and its content-selective pattern during *IS*. Left and right PPC were selected as memory ROIs and due to their being large and non-spherical clusters, we used the conjunction of the following contrasts to select all PPC voxels that showed significant effects: *IN*, *IN* > *IS*, *IN* MVPA and *IN* >*IS* MVPA. The resulting left PPC ROI entailed 120 voxels (centroid *x* = −20, *y* = −72, *z* = 40) and right PPC ROI entailed 548 voxels (centroid *x* = 24, *y* = −60, *z* = 54).

**Figure 3:**
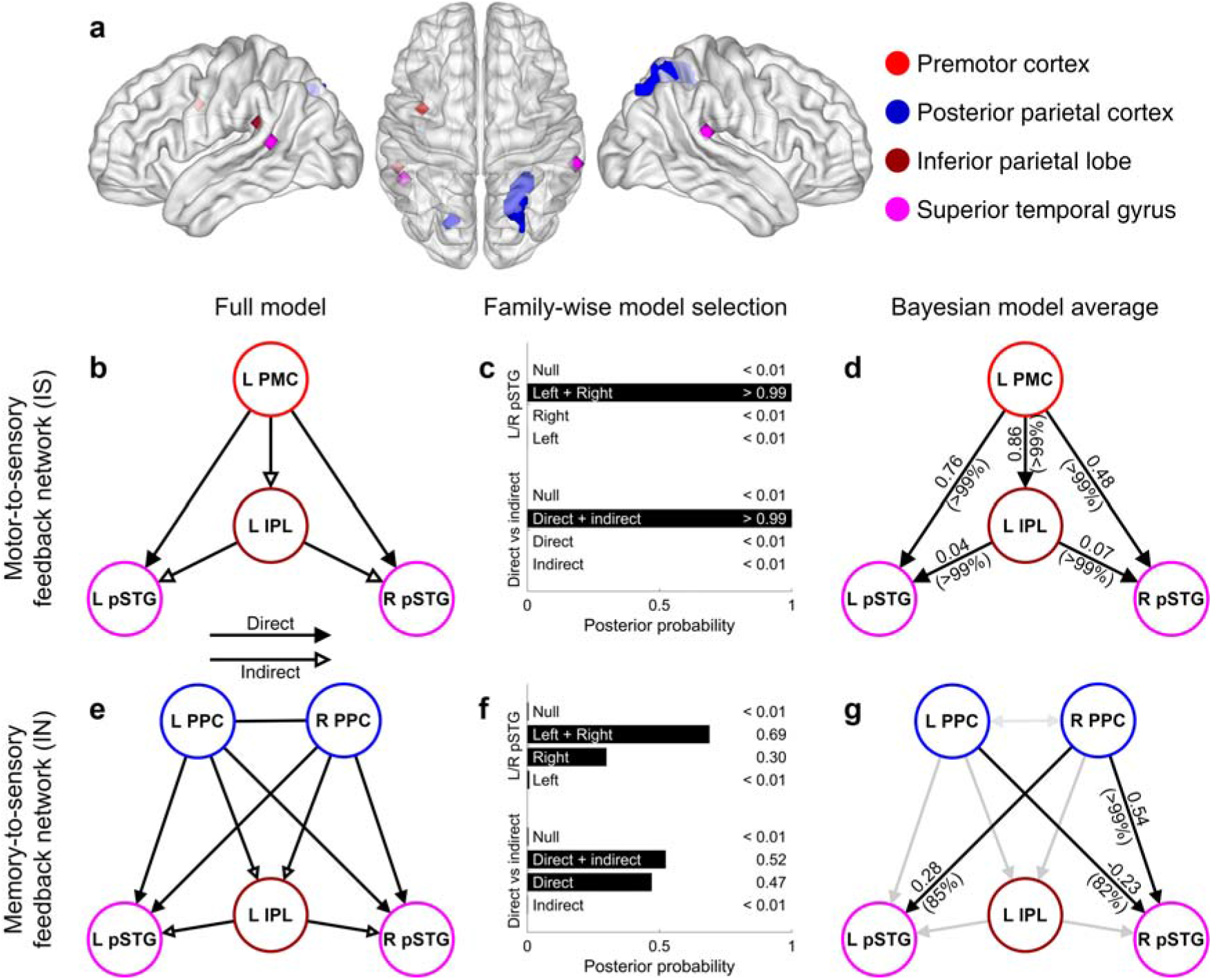
Motor-to-sensory and memory-to-sensory feedback networks assessed by dynamic causal modeling (DCM). (**a**)Regions of interest (ROIs) used for DCM. ROIs were spherical with a radius of 4 mm except for bilateral PPC ROIs, which were selected based on contrast conjunctions. (**b**) Graphical illustration of the full model of the motor-to-sensory feedback network. (**c**) Family-wise Bayesian model comparison of the two motor-to-sensory feedback model factors (feedback structure and sensory endpoints). Numbers on the right denote posterior probability. (**d**) Bayesian model average of effective connectivity parameters for the motor-to-sensory feedback network. Parameters that reached the significance level of posterior probability (Pp) > 0.75 were shown in black and otherwise in gray. Numbers out of paratheses denote parameter estimate in the unit of Hertz and numbers in paratheses denote posterior probability. (**e-g**) DCM results similar as (**b-d**) but for the memory-to-sensory feedback network.

To test our central hypothesis about the feedback projections from upstream motor and memory networks to the auditory cortex in generating content-specific prediction, we used dynamic causal modeling (DCM) (Friston et al., 2003), a well-established method that allows inference of directional brain connectivity modulated by an experimental condition (*IS* and *IN*, in the present study). DCM features a neuronal state equation which is coupled to a biophysically plausible model to explain BOLD signals (see Methods for details). Among all DCM parameters, our research question majorly concerns connectivity modulated by imagery. We specified and inverted two full motor-to-sensory and memory-to-sensory DCMs entailing all *a priori* imagery-modulated connectivity parameters ‘switched on’ using data from *IS*and *IN* sessions respectively.

To construct a full motor-to-sensory DCM, we allowed *IS* to modulate 5 connections: *direct* connections from left PMC to bilateral pSTG, and *indirect* connections from left PMC to left IPL and then to bilateral pSTG (Figure 3b). We then constructed 11 reduced models with a subset of these connections ‘switched off’ according to two factors, concerning the feedback architecture (direct-only / indirect-only / direct and indirect / null) and auditory endpoint (left pSTG / right pSTG / left and right pSTG / null). The null model contained no modulated feedback connection and thus offers the null hypothesis. A graphical illustration of all reduced models is shown in Figure S5.

Under the Bayesian Model Reduction (BMR) scheme (Friston et al., 2016; Zeidman et al., 2019a; Zeidman et al., 2019b), the free energy (lower bound on model evidence) (Friston et al., 2007) of each reduced model was derived. This allowed us to perform Bayesian model comparison (BMC) to systematically infer whether motor-to-sensory connections were enhanced in *IS* and if so, through what route (direct versus indirect) and in which hemisphere they ended. BMC returned the single winning model to be the full model itself with a posterior probability (Pp) higher than 0.99. We pooled reduced models according to the two factors to perform family-wise Bayesian model selection (Figure 3c), which revealed that the hybrid architecture entailing both direct and indirect connections to bilateral pSTG was the most likely (Pp > 0.99 for both families). We then summarized model parameters across all models by taking the weighted average of parameters from each model with the weight determined by each model’s Pp, an approach known as Bayesian model average (BMA) (Jennifer et al., 1999). The BMA results (Figure 3d) confirmed the essence of all 5 *IS*-modulated connections which all had a positive mean and Pp > 0.99. All these results together suggest a motor-to-sensory feedback architecture originating at the left PMC, mediated by left IPL. and ending at bilateral pSTG during *IS*.

A similar procedure was applied to construct and evaluate memory-to-sensory DCMs using data from the IN session. This DCM model specified entailed *IN*-modulated connections between bilateral PPC,*direct* connections from bilateral PPC to pSTG, and *indirect* connections from PPC to left IPL and then to bilateral pSTG (Figure 3e). 112 Reduced models were constructed according to 4 factors: feedback origin (left PPC / right PPC / left and right PPC), auditory endpoint (left pSTG / right pSTG / left and right pSTG / null), feedback architecture (direct-only / indirect-only / direct and indirect / null), and PPC mutual connection (present / absent).

BMC over reduced models of the memory-to-sensory DCM showed that the most probable (despite the relatively low Pp = 0.18) model entailed feedback connections initiating from bilateral PPC (without mutual connection) to bilateral pSTG via both direct and indirect pathways. Results of family-wise model selection over the two most important factors were shown in Figure 3f. Regarding the feedback architecture, evidence near equally supported the direct-only architecture (Pp = 0.47) and the hybrid architecture with both direct and indirect connections (Pp = 0.52). Bilateral PPC (Pp = 0.65) was more probable than right PPC alone (Pp = 0.35) to be the feedback source, and bilateral pSTG (Pp = 0.69) was more probable than right pSTG alone (Pp = 0.30) to be the auditory endpoints. When summarizing individual parameter estimates using BMA, we found three significant (Pp > 0.75) connections along the direct route (Figure 3g): left PPC to right pSTG (mean = −0.23 Hz, Pp = 0.82); right PPC to left pSTG (mean = 0.28 Hz, Pp = 0.85); and right PPC to right pSTG (mean = 0.54 Hz, Pp > 0.99). Overall, these results spoke for the existence of a memory-to-sensory projection from bilateral PPC to bilateral pSTG. *IN* modulated the left PPC to pSTG connection in an inhibitory manner while enhancing right PPC to pSTG connections, suggesting a hemispheric division of function. The lack of evidence in family-wise model selection and BMA did not support any mediating role of left IPL in memory-to-sensory feedbacks.

### Distinct motor-to-sensory and memory-to-sensory feedback networks in generating predictions

After mapping out the functional motor-to-sensory and memory-to-sensory feedback networks using *IS* and *IN* data respectively, we continued to ask whether the two networks were differentially implemented during *IS* and *IN*. To test our hypothesis of the two networks being functional distinct, we ‘swapped’ the data-model combination, that is, we re-inverted the full motor-to-sensory and memory-to-sensory DCMs (Figure 3b and Figure 3e) using data from the other imagery condition than that used in the previous section. That is, we used *IN* data (BOLD timeseries and imagery events in the condition) as inputs to the specified motor-to-sensory DCM and used *IS* data for memory-to-sensory DCM. We then compared variance explained by the DCM as well as PEB parameter estimates in each model fitted with *IS* and *IN* data.

We found that the motor-to-sensory DCM fitted with *IS* data yielded significantly higher explained variance (mean = 14.96%) than with IN data (mean = 6.86%), as revealed by a two-sided Wilcoxon-signed rank test (*P* = 0.01, Figure 4a). However, no significant difference in mean parameter estimates (*P* > 0.09 for all five parameters, two-sided *z*-test) was found (Figure 4b). These results suggest that the motor-to-sensory model cannot effectively explain *IN* data, despite the fact that ‘forced’ modeling fitting yielded similar parameter estimates.

**Figure 4:**
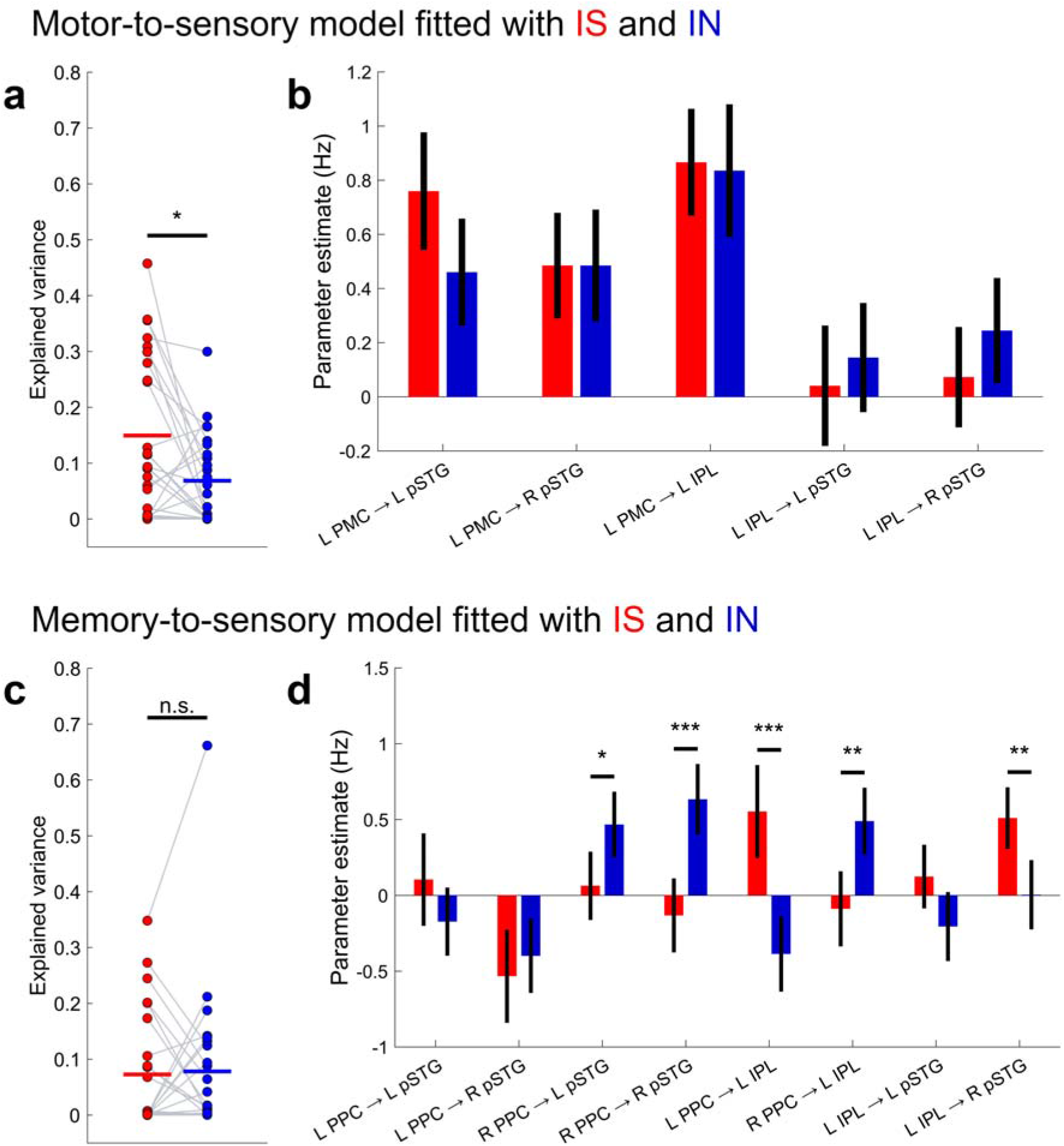
Motor-to-sensory and memory-to-sensory feedbacks differentially take place during IS and IN. (**a**) Variance explained by the motor-to-sensory DCM. Data for each individual are joined by a gray line. Red and blue denote results obtained with data from *IS* and *IN* sessions respectively. The means of *IS* and *IN* data are indicated by thick solid lines. (**b**)PEB estimates for all five imagery-modulated connections in the motor-to-sensory DCM. (**c-d**)Similar to (**a-b**) but for the memory-to-sensory DCM. Error bars indicate 95% confidence interval. \**P* < 0.05; \*\**P* < 0.01; \*\*\**P* < 0.001

On the other hand, we did not see a significant difference (*P* = 0.90) in explained variance when fitting the memory-to-sensory DCM with *IS* and *IN* data (mean explained variance = 7.28 and 7.82) (Figure 4c). Significant difference in several PEB parameters was observed (Figure 4d). Notably, the modulated connections from right PPC directly to bilateral pSTG were significantly higher in *IN* than in *IS* (right PPC to left pSTG, *P* = 0.033; right PPC to right pSTG, *P* < 0.001). Such differences suggest that the memory-to-sensory architecture identified in the previous section does not explain activities during motor-based prediction. Whereas several connections involving left IPL in the indirect pathway yielded higher parameter estimates using *IS* data (left PPC to left IPL, *P* < 0.001; left IPL to right pSTG, *P* = 0.006). These results were consistent with the indirect pathway found in the motor-to-sensory DCM, as left IPL exerted excitatory connectivity to pSTG even in a memory-to-sensory DCM where no motor node was included. These results also explained why there was no significant decrease in explained variance when fitting the memory-to-sensory DCM with *IS* data, as some pSTG activity might have been explained by IPL-exerted connectivity. In summary, the results in the analysis of swapping different types of imagery in fitting DCM suggest that distinct functional motor-to-sensory and memory-to-sensory feedback networks and different subregions of parietal lobe (IPL vs. PPC) mediate the generation of content-specific auditory prediction in *IS* and *IN*.

## Discussions

Using fMRI with a dual imagery paradigm, we have characterized the neural implementation of sensory prediction via feedback projections from motor and memory systems to the auditory cortex. Our results revealed the motor and memory systems as independent sources of prediction. The differential involvement of IPL and PPC in the motor-based and memory-based prediction pathways further suggests a functional division of the parietal lobe for routing the generation processes. The inter-areal communicative neural structures mediate distinct predictive processes via representational transformation, converging motor and memory information into sensory format for adaptive behavior.

### Motor-based prediction originates in the premotor cortex

Significant activity was observed in the left PMC in the motor-based prediction task of *IS* and its representational specificity was supported by MVPA (Figure 1,2). These results are consistent with previous studies that stress PMC’s role in speech planning (Castellucci et al., 2022) as well as studies on speech imagery (Li *et al*.,2020; Proix et al., 2022; Tian *et al*., 2016). In terms of lateralization, left PMC was more engaged in speech prediction. Crucially the directed connectivity from PMC to pSTG was enhanced by motor-based imagery, thus revealing PMC’s fundamental role as the upstream motor system in predicting the auditory consequence of speech. Two other common motor areas, preSMA and the inferior frontal gyrus (IFG, Brodmann area 44 and 45, see Figure S1) were also activated. Yet, neither of them possessed significantly decodable representations.

These results suggest that the efference copy, as previously hypothesized (Wolpert and Ghahramani, 2000), is transformed from a copy of the motor plan generated in PMC and sent to the auditory cortex in a feedback manner.

### Inferior parietal lobe relays motor-to-sensory predictive signaling

Motor-to-sensory information flow from the PMC to pSTG was achieved by both direct and indirect routes (Figure 3). The indirect route features IPL as a relaying hub (also referred to as the Sylvian parietal-temporal area, Spt). These results are consistent with previous reports of IPL activation in both speech perception and production (Buchsbaum et al., 2001; Hickok et al., 2003; Hickok et al., 2009).

The intermediate step of IPL in the motor-based prediction generation route could be an auditory-motor interface and computes the transformation between motor and auditory representations (Hickok, 2012). Alternatively, because movement of articulators yield speech, the computation of auditory prediction could be mediated by predicting the sensorimotor status of articulators (Tian and Poeppel, 2010; 2012). Thus, the IPL could be an intermediate stage for an abstract somatosensory prediction in a functional continuum between the somatosensory regions in anterior part of parietal lobe to the final auditory prediction starting in the posterior part of temporal lobe. Somatosensory prediction has been observed in secondary somatosensory area and extending to IPL (Kilteni and Ehrsson, 2020). In the speech domain, the partial redundant predictions in the sensorimotor and auditory domains may provide computational benefits of detecting distinct noise sources.

### Posterior parietal cortex mediates memory-to-sensory predictive signaling

PPC was active in the memory-based prediction task of *IN* and harbored imagery-specific codes in *IN* (Figure 1,2). DCM further revealed enhanced connectivity between right PPC and bilateral STG, suggesting right PPC is the crucial origin of the memory-to-sensory prediction network (Figure 3). The role of PPC in episodic memory has been demonstrated in a broad range of studies employing paradigms such as N-back (Barch et al., 2013; Owen et al., 2005), retention (Kwak and Curtis, 2022), and memory search (Sestieri et al., 2014). Directed connectivity from PCC to the sensory cortex has also been found in visual imagery (Dentico *et al*., 2014; Dijkstra *et al*., 2017). Altogether, these findings further support PPC as a general episodic buffer in generating memory-based prediction across memory tasks and modality.

Another interesting property is that left PPC to STG connectivity is reduced instead of enhanced as observed in its right PPC to STG counterpart. This could be due to a hemispheric division of PPC in auditory memory or a functional-anatomical division of PPC, as the left PPC ROI we selected is majorly composed of the intraparietal sulcus while the right PPC ROI majorly consists of the superior parietal lobule.

The prefrontal cortex and hippocampus were less supported by empirical evidence to be the origin in the memory-to-sensory network as they lacked significantly decodable patterns. As the role of vlPFC and hippocampus in memory maintenance and memory-based prediction has been described in the literature (Davachi and DuBrow, 2015; Kumar et al., 2016), the discrepancy may arise from the experimental design and analysis scheme. Throughout our analyses, we modeled the imagery events as sustained boxcar events. Since participants may recall the soundtrack of the videos immediately after their initiation appearance, vlPFC and hippocampus could support the initial retrieval of auditory memory through visual-auditory association which is then transferred to PPC for maintenance. The interpretation is however hard to assess due to the low temporal resolution of fMRI.

Outside of DSPM, we also found that the cingulo-opercular network (CON, including FO/aINS and dACC/dmPFC) was more active in *IN* but lacked decodable multivoxel patterns. This is consistent with previous studies reporting CON to have a more modulatory rather than representational role in memory (Sestieri *et al*., 2014; Wallis et al., 2015). Because our study focuses on representational transformations in feedback projections, we did not include CON in DCM to avoid complicating the model. Yet our data suggest CON may have a role in modulating memory-based prediction and imagery.

### Common auditory reactivation via different feedback projections

Common activation in both motor-based and memory-based imagery in the auditory cortex agrees with previous work on musical imagery (Halpern and Zatorre, 1999; Li *et al*., 2020), speech imagery (Proix *et al*., 2022; Tian *et al*., 2016), and imagery of complex sounds (Bunzeck *et al*., 2005). Imagery induced similar activations in the auditory cortices as hearing controls, supporting the nature of sensory-like representation as the ending result of prediction. The commonality in auditory reactivation in *IS* and *IN* further suggests a sensory convergence of predictions originating in different upstream networks. Together, the common sensory activations by hearing and types of imagery hint at a neuroanatomical foundation for the integration of various predictive and stimulus-driven signals in the sensory system. At a more microscopic level, such integration may take place in distinct neural subpopulations in the sensory system that differentially respond to feedforward sensation and feedback prediction. A likely laminar organization for such functional populations involves feedback prediction sent to the deep layers of the sensory cortex (Kok et al., 2016; Rao and Ballard, 1999). Further investigations adapting our paradigm can aim to test this specific hypothesis, which will shed light on the microcircuitry that integrates feedforward input and feedback prediction.

### Conclusions

In conclusion, using a dual imagery paradigm with fMRI, we found that motor and memory systems project to the sensory system via distinct network structures to generate sensory predictions. The neural origin and inter-areal communicative structures constrain the computations of representational transformation, creating the emergent properties of the distinct predictive neural networks for efficiently linking cognition with environment.

## Methods

### Ethics statement

The experimental protocol was approved by the Institutional Review Board at New York University Shanghai (IRB00009975/FWA#00022531) in accordance with policies and regulations found in The Common Rule (45 CFR part 46).

### Participants

Twenty-nine right-handed native Mandarin speakers participated in the experiment with informed consent and received monetary incentives. No participant reported a history of neurological or psychological illness. All participants had normal or corrected-to-normal vision. Data from four participants were removed from analyses due to excessive head motion or drowsiness during scanning. The remaining 25 participants were included in the analyses (12 females; mean age ± standard deviation = 21.3 ± 2.3).

### Materials

Ten different seven-second video clips with their corresponding audio tracks were selected and used as the stimuli in the experiment. All video clips were about scenes or objects and none of them contained human speech. Examples included a basketball bouncing on the wooden floor, a training quickly passing by, and a ringing telephone. Our motivation was to choose videos with sounds that were hard to simulate with human vocal organs, but easy to imagine with the aid of visual scenes. Every 500 ms a square image patch was superimposed on the center of the video, making a total of 14 patches. These images were either Chinese characters (black, against a white background) constituting a sentence that described the content (see Table S1) of the video clip or mosaics made by randomly shuffling pixels of the original character image, thus ensuring equal net luminance as the character images. We also created synthesized speech of the sentences in a male’s voice using the VoiceGen toolbox (https://github.com/ray306/VoiceGen).(https://github.com/ray306/VoiceGen).

### Procedure

We presented participants with 12 sessions of videos following a structural scanning session. Trials in every session shared a similar procedure: fixation period (500 ms), video presentation (7 s), vividness rating/catch trial detection (< 3 s), and an inter-trial interval of either 4.44 or 6.66 s (2 or 3 repetition time for fMRI scanning) minus the response time for rating or detection task. If a participant did not press any button within 3 seconds or reported incorrectly during catch trial detection, the trial was considered as a no-response or wrong-response trial that was separately modeled, and thus excluded from further analyses.

During the first 3 sessions, participants were presented with videos with the original audio tracks with mosaics overlaid on them. We refer to this condition as *Hearing of Non-speech sounds (HN*). Each HN session consisted of 22 trials which included two catch trials featuring a pure tone (frequency = 1000 Hz, duration = 715 ms) played at a random time point of a random video. The other 20 trials consisted of ten videos each played twice in random order. After watching each video, participants were asked to report if they heard the pure tone in the video by pressing button 1 (for yes) or button 2 (for no) on an MRI-compatible response pad.

Three sessions of *Imagery of Non-speech (IN*) followed. In these sessions, videos were muted, and mosaics were overlaid in the center. Participants were instructed to imagine the sounds they heard during the preceding *HN* sessions and rated the vividness of imagery (rating range = 1 - 5) with the response pad. This visually aided imagery of the non-speech task was similar to previous studies (Bunzeck *et al*., 2005). Thereafter came three sessions of *Imagery of Speech (IS*)where the videos were also muted, and Chinese characters were overlaid on the videos. Participants were instructed to imagine saying the characters and gave a vividness rating afterwards. Every *IS* or *IN* session consisted of 20 videos with each of the 10 videos randomly played twice.

The task in the last three sessions was *Hearing of Speech (HS*). During the video presentation, the original audio track was replaced with synthesized speech. Similar to the *IN* sessions, 2 catch trials were included in each session in which two nearby characters in the synthesized speech were reversed (e.g., 鞭炮 to 炮鞭; firecracker to ‘crackerfire’). Participants indicated whether they heard a reversal using the response pad in a similar manner as in *HN* sessions.

The hearing sessions (*HN* and *HS*) were designed to localize areas that are activated by external auditory stimuli in a content-specific manner. The order of sessions (*HN-IN-IS-HS*) was designed such that participants were first familiarized with original sounds in the videos during *HN*, and were able to recall them during *IN*. *IN* sessions proceeded *IS* and *HS* sessions such that participants were unaware of and less likely to perform imagery of speech during *IN*. *IS* proceeded *HS* because there otherwise existed an alternative strategy for participants to retrieve their memory of the synthesized speech they had listened to.

### fMRI data acquisition

MRI images were collected on a Siemens MAGNETOM Prisma^fit^ System (Erlangen, Germany) at East China Normal University. Anatomical images were acquired using a T1-weighted magnetization-prepared rapid acquisition gradient echo (MP-RAGE) sequence (192 sagittal slices; field of view (FOV) = 240 mm × 240 mm; flip angle (FA) = 8°; repetition time (TR) = 2300 ms; echo time (TE) = 2320 ms; voxel size = 0.9375 × 0.9375 × 0.9000 mm^3^). Functional images were acquired using a T2*-weighted echo-planar imaging (EPI) pulse sequence (38 even-first interleaved slices; FOV = 192 mm × 192 mm; FA = 81°; TR/TE = 2220/30 ms; voxel size = 3.0 × 3.0 × 3.6 mm^3^; interslice gap = 0.6 mm). Functional slices were oriented to an approximately 30° tilt toward coronal from AC–PC alignment to maximize coverage of individual brain volumes.

### Preprocessing

Preprocessing of fMRI data and subsequent analyses were implemented via SPM12 (https://www.fil.ion.ucl.ac.uk/spm/, version 7771) and custom-written scripts with MATLAB R2021a (MathWorks Inc., Natick, MA, USA). Preprocessing followed the standard procedure in SPM12.

All functional images from each participant were temporally interpolated to the first slice of each volume and spatially realigned to the mean image. The structural image was co-registered with functional images. For univariate and DCM analyses, functional images were then spatially normalized to the Montreal Neurological Institute (MNI) standard brain space (resampled voxel size = 2 mm isotropic) and smoothed with a 6 mm full width, half maximum (FWHM) Gaussian kernel. For MVPA, the functional images were neither normalized nor smoothed to preserve information patterns in the individual’s native brain space.

### Univariate analysis

Events were modeled as sustained boxcar epochs spanning their corresponding duration. They included the presentation of fixation points, videos (in which participants performed the imagery and hearing tasks), instructions for vividness rating or catch trial detection, and button presses. Events in catch trials, no-response, or wrong-response trials were modeled separately to improve model sensitivity. All events were convolved with a canonical hemodynamic response function (HRF) implemented in SPM12 and entered as regressors into a general linear model (GLM) for each individual. Each GLM also included head motion regressors and session-wise baseline regressors. The GLM was then estimated using functional data high pass filtered at 1/128 Hz. Individual-level contrasts were constructed using the beta estimates of regressors of interest and were subject to a one-sample *t*-test for group-level inference.

To examine common activation in imagery and comparable hearing conditions (Figure S2), we took a minimum statistics approach (Nichols *et al*., 2005). We first obtained thresholded *t*-value-maps from imagery and hearing conditions (e.g.,*IS* and *HS*), computed the minimum *t*-value from the two conditions for each voxel, and reported only voxels that were significant after the operation (*t*_24_ > 2.80,*P* < 0.005). Similarly, to examine differential activations in *IS* and *IN* in Figure 1d, we took the minimum *t*-value from the *IS* and *IS* > *IN* contrasts as well as *IN* and *IN* >*IS* contrasts. Therefore, all significant voxels revealed had both significant activities during one type of imagery and significant difference over the other.

### Multivoxel pattern analysis (MVPA)

One additional participant was excluded from MVPA due to his lack of response to the coin video in all *HS* sessions, making the sample size *N* = 24. MVPA was conducted using The Decoding Toolbox (TDT, version 3.999E) (Hebart et al., 2015). Beta estimates of each video category from all three sessions in imagery (IS and IN) or hearing (*HS* and *HN*) conditions were obtained and were used to train and test a L2-norm support vector machine (SVM) available through LIBSVM (Chang and Lin, 2011). We used a regularization parameter C = 1 and scaled the data at a range of 0 to 1. To efficiently test which voxels across the brain could be used for accurate classification, we moved spherical searchlights (Kriegeskorte et al., 2006) throughout the brain. To avoid the choice of radius from biasing our results, we conducted searchlight analyses with varying radii from 1 - 8 voxels. The accuracy maps obtained using a 4-voxel radius were visualized as surface renderings.

To decode video categories within a condition (Figure 2; Figure S3), we used a leave-one-session-out cross-validation scheme. In each decoding step, two out of three sessions in the condition were used to train an SVM classifier and the remaining session was used as test data to decode the 10 video categories from multivoxel patterns. The average classification accuracy from all 3 decoding steps, each having a different test session and two corresponding training sessions, was calculated and assigned to the center voxel of the searchlight to generate a decoding accuracy map.

To decode video categories across *IS* and *IN* data (Figure S4), we used a two-way leave-one-session-out cross-classification scheme. Similar to the previous scheme, two sessions from the *IS* condition were used to train a classifier, but the test data this time was one session from the *IN* condition. This procedure was iterated for all three *IN* sessions, each using a different combination of two *IS* sessions as training data. Next, the *IS* sessions were used as test data for classifiers trained with *IN* sessions. A cross-classification accuracy map was generated using the average accuracy obtained from a total of 6 decoding steps.

To perform group-level inference, we normalized the individual-level accuracy maps into MNI space and smoothed them with a 6 mm FWHM Gaussian kernel to account for the individual neuroanatomical difference. These accuracy maps were then brought to one-sample *t*-tests and the mean accuracy level of each significant cluster (voxel-wise threshold: *P* < 0.005; cluster-wise threshold: *P*_FDR_ < 0.05) was displayed. For better visualization, the range of data for display was controlled at 10 – 20% (0 – 10% above the chance level of 10%) because accuracy above 20% was mostly observed in visual areas.

### Timeseries extraction from regions of interest (ROIs)

Two types of ROIs were used for effective connectivity analyses. Spherical ROIs (left and right pSTG, left IPL, and left PMC) consisted of gray matter voxels within spheres whose center coordinates have been reported in Table S2. The radius of each sphere was 4 mm. Specifically, this small radius ensured that the left pSTG and left IPL ROIs that were situated nearby (Euclidean distance = 14.97 mm) had no shared voxels nor smoothing-induced (FWHM = 6 mm) data contamination. On the other hand, since gray matter voxels in bilateral PPC that were significant in univariate and MVPA analyses were clearly non-spherical, we created a mask using all significant voxels in bilateral PPC and used it to select the left and right PPC voxels.

For each ROI, voxel-wise time courses in *IS* and *IN* were high pass filtered at 1/128 Hz and the estimated effects of non-imagery regressors (e.g., fixation cue, button press, and head motion) were subtracted out. This adjustment should increase DCM model sensitivity by excluding activities induced by non-imagery events. The resulting first principal component of each ROI was used for DCM analyses.

### Dynamic causal modeling (DCM)

We used the bilinear DCM that features the following state equation:

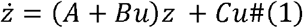

where *z* denotes hidden neural activity from all ROIs and the dot notation denotes change per unit time. The *A* matrix represents baseline connectivity in the absence of external stimulation. The *B* matrix represents the modulatory effects of an experimental input *u (IS* or *IN* in the present study) on connectivity between regions. The *C* matrix represents the direct driving effect of each *u* on neuronal activity.

Since we are interested in imagery-modulated connectivity, *B* matrix parameters are the most crucial to our study and those enabled connections for each model have been described in the main text. Regarding the rest of the parameters, we specified an all-ones A matrix (i.e., enabled baseline connectivity between every ROI pair) for both motor-to-sensory and memory-to-sensory models because we did not have any prior hypothesis regarding baseline connectivity. In terms of driving inputs, we specified imagery conditions to drive the motor and memory networks identified in univariate and MVPA analyses. We set *IS* as the driving input to left PMC in the motor-to-sensory model and *IN* as the driving input to bilateral PPC in the memory-to-sensory model. Enabled parameters had Gaussian priors with zero mean and non-zero variance while the others had zero variance.

The neural activity *z* was coupled with a biophysically informed forward model (Friston *et al*., 2003; Zeidman *et al*., 2019a) to predict the BOLD timeseries. The default one-state, stochasticity-not-included version of DCM was used. A slice timing model was used in alignment with the slice timing correction performed during preprocessing.

For subject-level model inversion, our goal was to find parameter estimates that maximize log model evidence. DCM uses a variational Laplace scheme to approximate model evidence with negative variational free energy (Friston *et al*.,2007). This estimation scheme also penalizes model complexity calculated as the Kullback-Leibler divergence between the priors and the posteriors. Thus, DCM evaluates how well the model achieves a trade-off between accuracy and complexity.

The expected parameter values and their covariance matrices at the subject level were then brought to a Parametric Empirical Bayes (PEB) analysis to make inferences about group-level effects (Friston *et al*., 2016; Zeidman *et al*., 2019a; Zeidman *et al*., 2019b). In terms of the between-subject design matrix, since our experimental design involves no between-subject factors, the design matrix was simply an all-ones vector *X =* [1 1…1 1]^T^ to model commonalities across subjects. In addition, random effects (unexplained between-subject variability) on parameters were assumed to account for individual differences.

Having estimated parameters of the motor-to-sensory and memory-to-sensory full models and specified candidate reduced models by ‘switching off’ some parameters, we then performed Bayesian Model Reduction (BMR) (Friston *et al*.,2016) to analytically derive the evidence and parameters of the reduced models. We compared the evidence of each reduced model to find the winning model described in the main text as well as pooled evidence of models belonging to each model family (Figures S5, S6). In Figure 3b and Figure 3, we plotted parameters that had positive evidence (Pp > 0.75) of being present vs absent, assessed by Bayesian Model Average (BMA) on all reduced models.

For data-model swapping in Figure 3d-g, we used the full model structures as in Figure 3b-c and fitted them with *IS* and *IN* data. The explained variance for each data-model combination was plotted in Figure 3d&f and two-sided Wilcoxon signed-rank tests were performed. The estimated group-level parameters were plotted in Figure 4e&g. Because the parameter estimates corresponded to a multivariate Gaussian density, we plotted the means and 95% confidence intervals computed with the leading diagonal of the covariance matrix. To compare the distributions yielded by a model with different data, we performed *z*-tests using the mean and variance of the parameter estimates.

## Contributions

O.M. and X.T. designed the experiment. O.M. collected the data. Q.C., O.M., and Y.H. performed the analyses. Q.C., O.M., and Y.H. drafted the manuscript. All the authors reviewed and corrected the manuscript. X.T. supervised the project.

## Acknowledgements

We thank Zheng Li, Xiao Ma, Wenjia Zhang, Yidan Gao, Lechuan Wang, Hao Zhu, Rui Tong, and Jialin Chen for their help in experimental design, data collection and analysis.

This study was supported by the National Natural Science Foundation of China 32071099, Natural Science Foundation of Shanghai 20ZR1472100, Program of Introducing Talents of Discipline to Universities, Base B16018, and NYU Shanghai Boost Fund to XT, and NYU Shanghai Dean’s Undergraduate Research Fund to Q.C.

## Code and Data Availability

All relevant data (including preprocessed functional images, event files, statistical maps, and dynamic causal models) and codes for replicating the key findings are available at https://doi.org/10.17605/OSF.IO/7492E.

