## Supplementary Information for "Dual-stream cortical feedbacks mediate sensory prediction"

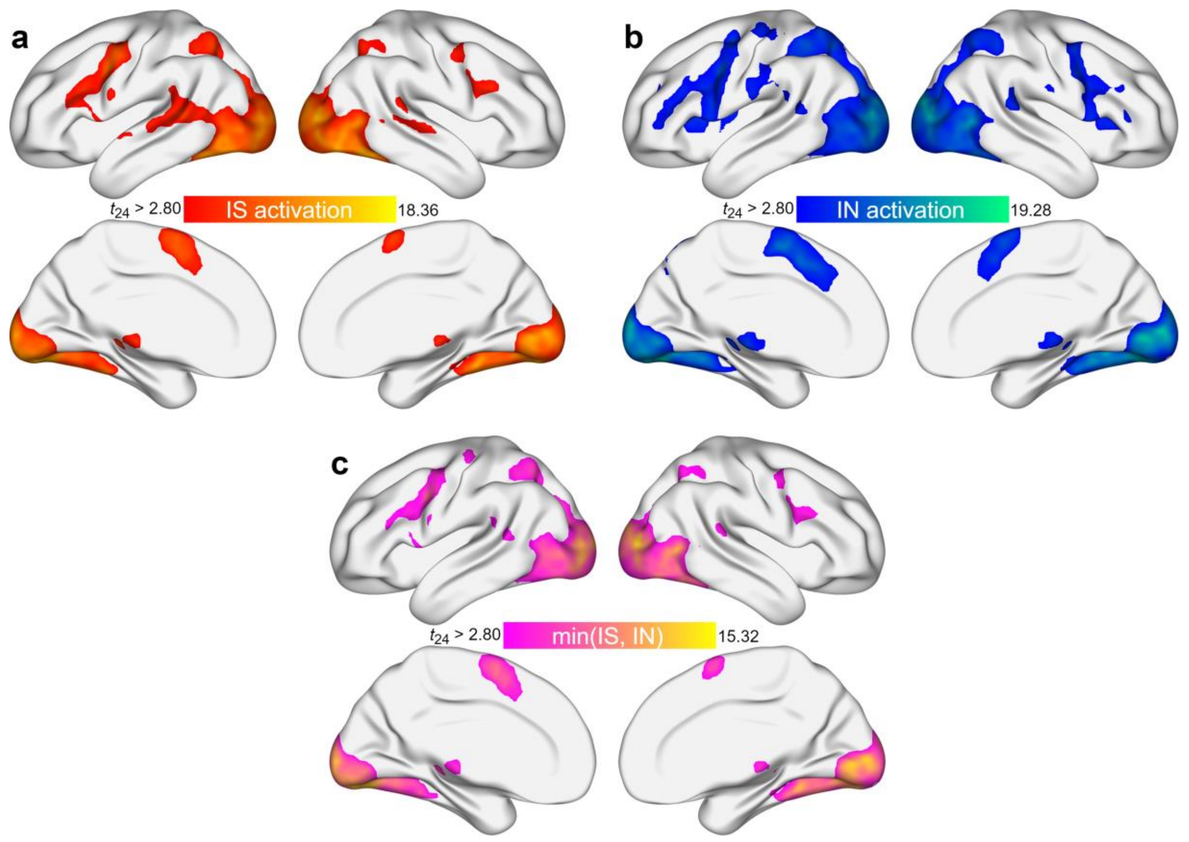


**Figure S1: Whole-brain activations in two imagery conditions and their conjunction, related to Figure 1.**

**(a)** Imagery of Speech (*IS*) **(b)** Imagery of Non-speech (*IN*) **(c)** The minimal stats conjunction between IS and IN. All maps were thresholded at $P < 0.005$ voxel-wise and $P_{FDR} < 0.05$ cluster-wise.

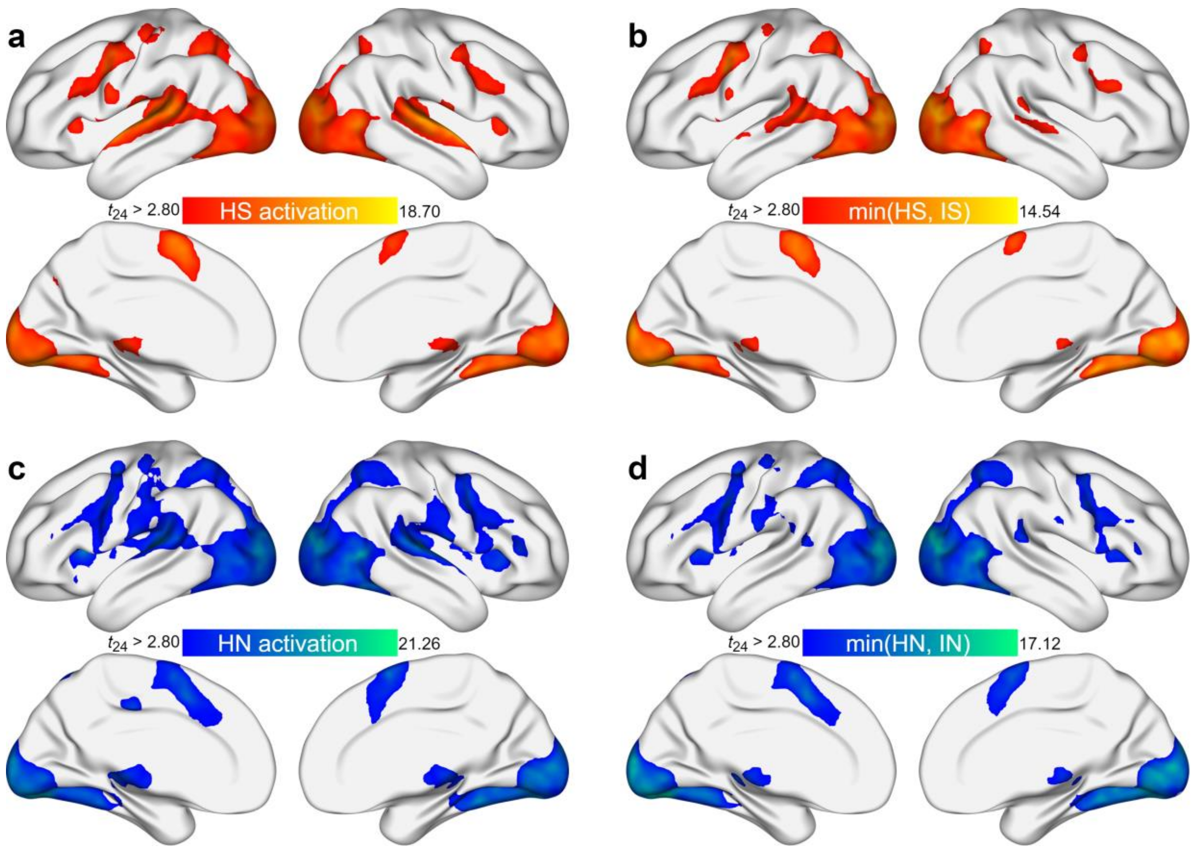


**Figure S2: Whole-brain activations in two comparable hearing conditions and their conjunction.**

**(a)** Hearing of Speech (*HS*), during which participants listened to synthesized speech of the sentence. **(b)** The minimal stats conjunction between *HS* and *IS*. Overlapped activation over STG and IPL suggests the processing of feedforward and feedback signals take place at same cortical sites. **(c)** Hearing of Non-speech sounds (*HN*), during which participants watched the video with original soundtracks. **(d)** The minimal stats conjunction between *HN* and *IN*. Similar to **(b)**, overlap over STG and IPL was found. All maps were thresholded at $P < 0.005$ voxel-wise and $P_{FDR} < 0.05$ cluster-wise.

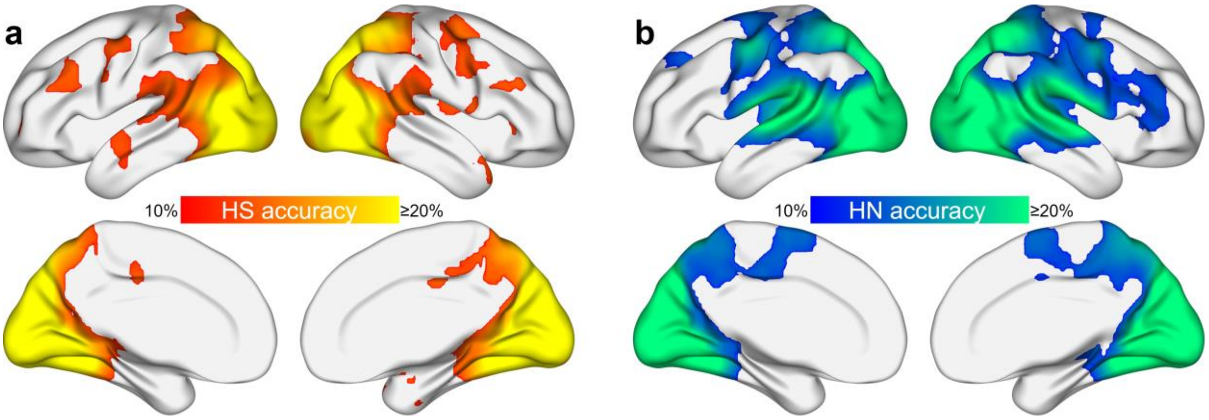


**Figure S3. MVPA results of hearing conditions**

**(a)** Hearing of Speech (HS). **(b)** Hearing of Non-speech sounds (*HN*). Accuracy maps were thresholded at $P < 0.005$ voxel-wise and $P_{FDR} < 0.05$ cluster-wise.

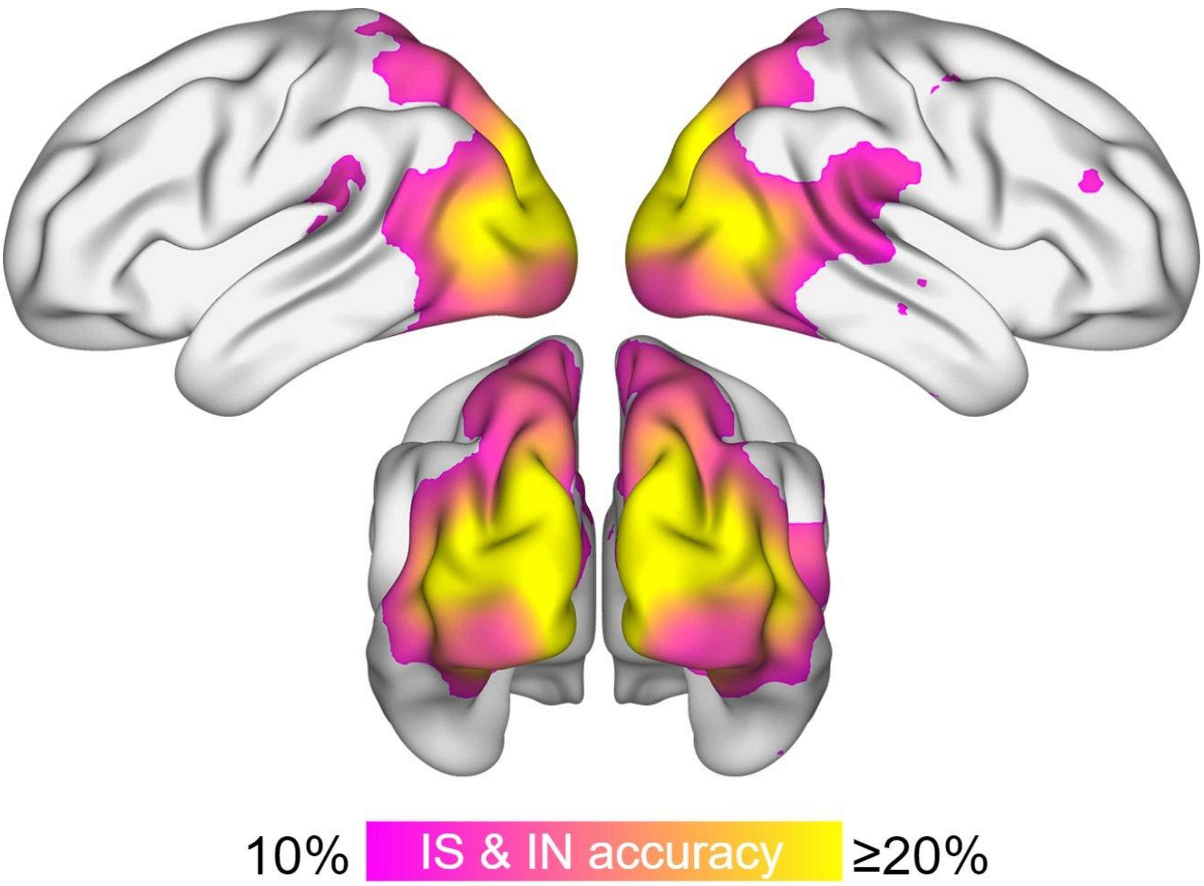


**Figure S4. Cross-classification of IS and IN reveals representational similarity in bilateral PPC.** The accuracy map was thresholded at $P < 0.005$ voxel-wise and $P_{FDR} < 0.05$ cluster-wise.

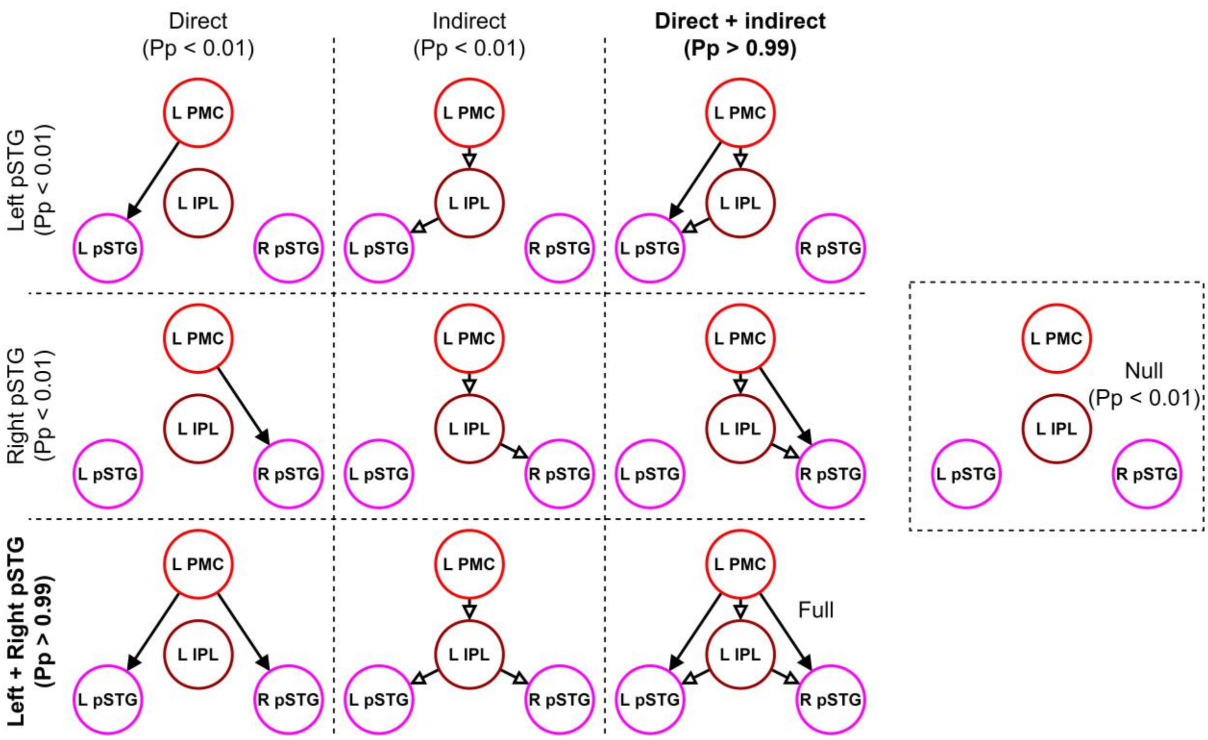


**Figure S5. Candidate models for the motor-to-sensory DCM.**

Enabled *IS*-modulated connections between ROIs are illustrated. Filled arrows denote direct connections from PMC to pSTG and unfilled arrows denote connections along the indirect route. Each row represents a family of auditory endpoint (left and/or right pSTG). Each column represents a family of feedback architecture (direct-only, indirect-only, and direct and indirect). The null model contains no imagery-modulated connections. Pp: posterior probability for the model family.

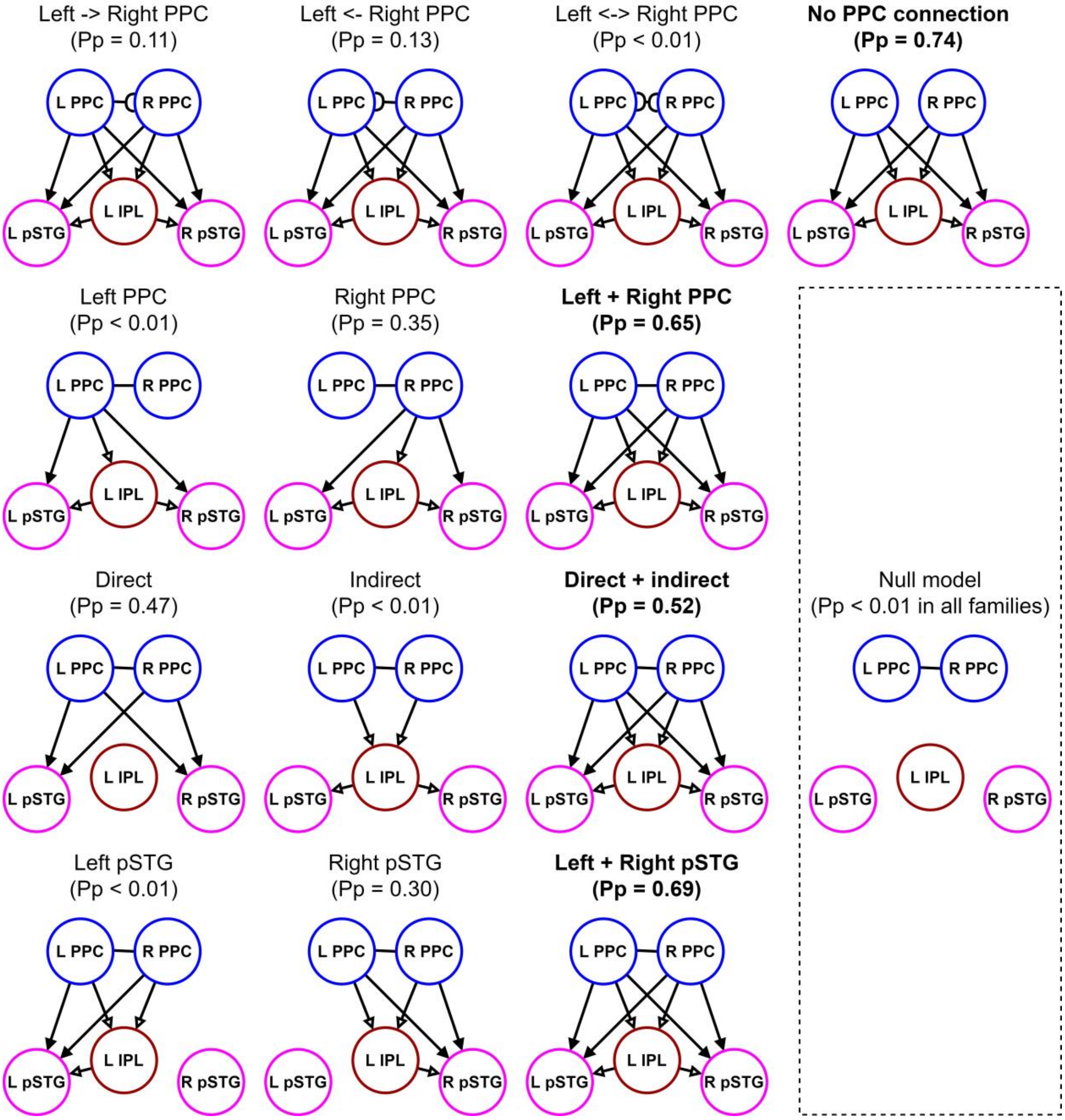


**Figure S6. Candidate models for the memory-to-sensory DCM.**

Similar to Supplementary Figure 5 but for *IN*-modulated memory-to-sensory connections. A total of 112 models were specified for the memory-to-sensory DCM according to 4 factors represented in each row. A complete graphical illustration of all 112 models would be redundant, therefore we illustrate here the representative models in each family with key connection(s) removed from the full model. The null model has no modulated connections except possible connections between bilateral PPCs. Pp: posterior probability for the model family.

| English translation |
| --- |
| Red firecrackers are popping off in the yard. |
| The second hand of a clock is rotating non-stop. |
| Water in the toilet is spinning relentlessly. |
| A basketball is bouncing on the wooden floor over and over. |
| A blue train is rapidly passing by. |
| A white helicopter is taking off in the airport. |
| Thunder roars as the lightning tears the sky apart. |
| A handful of coins are dropped on the hard floor. |
| A transparent stream is gently flowing through the stones. |
| The old-style Nokia phone is vibrating as a call comes in. |

**Table S1: The list of Chinese sentences used in IS and HS sessions.**

| Anatomical location | Center coordinate | | | Center *t* value | | | | | | |
|---|---|---|---|---|---|---|---|---|---|---|
| | | | | Univariate | | | | MVPA (radius = 4) | | |
| | x | y | z | IS | IS > IN | IN | IN > IS | IS | IN | IN > IS |
| L (Posterior) Superior temporal gyrus (L pSTG) | -50 | -46 | 12 | 5.36 | n.s. | 4.43 | n.s. | 4.40 | 5.16 | n.s. |
| R (Posterior) Superior temporal gyrus (R pSTG) | 62 | -36 | 18 | 5.64 | n.s. | 6.24 | n.s. | 3.24 | 4.81 | n.s. |
| L Inferior parietal lobe (L IPL) | -54 | -38 | 24 | 4.37 | n.s. | 2.90 | n.s. | 4.28 | 3.76 | n.s. |
| L Premotor cortex (L PMC) | -38 | 0 | 36 | 6.76 | 3.13 | 4.99 | n.s. | 3.86 | 3.74 | n.s. |
| L Posterior parietal cortex (L PPC) | -20 | -72 | 40 | n.s. | n.s. | 5.74 | 4.48 | 5.41 | 7.00 | 3.46 |
| R Posterior parietal cortex (R PPC) | 24 | -60 | 54 | n.s. | n.s. | 6.88 | 3.96 | 5.09 | 5.53 | 3.54 |

**Table S2**: Selected regions of interest for DCM analysis. Left and right pSTG, left IPL and left PMC ROIs were specified as spheres with a radius of 4mm. The center coordinates (in MNI) are provided in the table. Each ROI comprised around 33 voxels. Left and right PPC ROIs, marked with an asterisk, were not spherical as they were selected by univariate and MVPA contrast conjunctions. The voxels closest to their geometric centers are reported here. $P < 0.005$ was considered not significant (n.s.).
